# *Vibrio cholerae* biofilm dispersal regulator causes cell release from matrix through type IV pilus retraction

**DOI:** 10.1101/2021.05.02.442311

**Authors:** Praveen K. Singh, Daniel K.H. Rode, Pauline Buffard, Kazuki Nosho, Miriam Bayer, Hannah Jeckel, Eric Jelli, Konstantin Neuhaus, Eva Jiménez-Siebert, Nikolai Peschek, Timo Glatter, Kai Papenfort, Knut Drescher

**Affiliations:** Max Planck Institute for Terrestrial Microbiology, Karl-von-Frisch-Str. 16, 35043 Marburg, Germany; Department of Physics, Philipps-Universität Marburg, Karl-von-Frisch-Str. 16, 35043 Marburg, Germany; Institute of Microbiology, Friedrich Schiller University, 07745 Jena, Germany; Biozentrum, University of Basel, Spitalstrasse 41, 4056 Basel, Switzerland

## Abstract

The extracellular matrix is a defining feature of bacterial biofilms and provides structural stability to the community by binding cells to the surface and to each other. Transitions between bacterial biofilm initiation, growth, and dispersion require different regulatory programs, all of which result in modifications to the extracellular matrix composition, abundance, or functionality. However, the mechanisms by which individual cells in biofilms disengage from the matrix to enable their departure during biofilm dispersal are unclear. Here, we investigated active biofilm dispersal of *Vibrio cholerae* during nutrient starvation, resulting in the discovery of the conserved *Vibrio* biofilm dispersal regulator VbdR. We show that VbdR triggers biofilm dispersal by controlling cellular release from the biofilm matrix, which is achieved by inducing the retraction of the mannose-sensitive hemagglutinin (MSHA) type IV pili and the expression of a matrix protease IvaP. We further show that MSHA pili have numerous binding partners in the matrix and that the joint effect of MSHA pilus retraction and IvaP activity is necessary and sufficient for causing biofilm dispersal. These results highlight the crucial role of type IV pilus dynamics during biofilm dispersal and provide a new target for controlling *V. cholerae* biofilm abundance through the induction and manipulation of biofilm dispersal.

## Introduction

Bacteria commonly live in biofilm communities, where cells are embedded and immobilized within a self-produced extracellular matrix (1, 2). Biofilms are difficult to eradicate in medical settings and technological flow systems (3–5), which can result in a reservoir for chronic bacterial contamination (6, 7). Although cells in biofilms often display a lower single-cell growth rate than planktonic cells, biofilm formation can provide important fitness benefits to the constituent cells (8–10). These beneficial community properties are emergent functions of biofilms, which include an increased tolerance to abiotic stresses, as well as increased protection from phage predation, social cheaters, and population invasion (8, 11–13).

Despite the fitness advantages that biofilms provide for bacterial cells, it can also be beneficial for cells to disperse from biofilms in adverse conditions, to colonize new habitats (14–19). However, biofilm dispersal is not a trivial process, as the cells are anchored in biofilms through bonds with the extracellular matrix. For cells to disperse from biofilms, it is therefore necessary that either the matrix is disintegrated or the bonds between the cells and the matrix are broken, before cells can actively depart from biofilms using flagellar motility, or passively depart *via* diffusion. How exactly cells are liberated from the biofilm matrix is generally unclear.

Fortunately, the matrix composition has been characterized for several biofilm model organisms over the last two decades, which paves the way for mechanistic studies of biofilm dispersal. For *V. cholerae*, the biofilm matrix primarily consists of the long-chain polysaccharide VPS, which is bound by the major matrix protein RbmA (20–22). Further matrix components are extracellular DNA (23, 24), the RbmC protein and several additional matrix proteins whose functions are mostly uncharacterized (25, 26), as well as factors used for surface attachment of the cells, including Bap1, FraH, CraA, and MSHA pili (20, 27–32). During *V. cholerae* biofilm growth, RbmA is known to be processed by the proteases HapA, PrtV, and IvaP (33, 34), yet neither of these proteases is known to play a role in dispersal from abiotic surfaces (35). Lack of the LapG protease, which cleaves FraH and CraA (27), can cause a dispersal defect in a static biofilm growth model for *V. cholerae* (36). Deletions of the extracellular nucleases Xds and Dns, which modulate extracellular DNA levels in biofilms, also cause a dispersal defect in static growth conditions (23). The putative polysaccharide lyase RbmB has been shown to reduce cell density inside biofilms (37), and deletion of *rbmB* causes VPS accumulation (38), as well as a reduction of biofilm dispersal in a static biofilm model (36). However, neither of the factors that have been implicated in biofilm dispersal defects have been shown to be sufficient for causing biofilm dispersal. Therefore, the precise mechanisms by which cells liberate themselves from the matrix during active biofilm dispersal remain unknown.

To investigate biofilm dispersal mechanisms of *V. cholerae*, we previously developed an assay to reliably trigger biofilm dispersal of *V. cholerae* in flow chambers, which is achieved by removing the carbon source from the inflowing medium, or by stopping the flow in the channel (39, 40). This assay was used to identify that the quorum sensing master regulator HapR and the general stress response sigma factor RpoS jointly control biofilm dispersal (39). Using this flow chamber assay, we now investigated how individual *V. cholerae* cells actively disengage themselves from the biofilm matrix during dispersal. We discovered a *Vibrio* biofilm dispersal regulator and an unforeseen mechanism for cell release during biofilm dispersal, based on the combination of type IV pilus retraction and processing of a particular matrix component, which is both necessary and sufficient for *V. cholerae* biofilm dispersal.

## Results

In chambers with a continuous flow of minimal M9 medium that contains glucose as the sole carbon source, *V. cholerae* biofilm growth is highly reproducible (37, 39, 41). Under these conditions, biofilm dispersal can be triggered by switching the feeding flow to M9 medium that lacks the carbon source (39). When biofilm dispersal is tracked at single-cell resolution, we observed that the local cell density inside the biofilm gradually reduces and the whole biofilm biovolume shrinks (Fig. 1A). We used this combination of 3D confocal microscopy and automated image analysis to determine the biovolume change of mature biofilms to quantify biofilm dispersal for numerous different conditions in this study. To determine if the biovolume change during carbon starvation is caused by *de novo* synthesis of dispersal effectors, or whether it is caused by the activation of premade effectors, we measured the biofilm biovolume change following glucose removal in the presence or absence of a translation inhibitor (chloramphenicol) and a transcription inhibitor (rifampicin). These measurements showed that if transcription and/or translation are inhibited, biofilm dispersal is strongly reduced (Fig. 1B), indicating that the underlying molecular processes rely on the synthesis of new transcripts and proteins.

**Figure 1.**
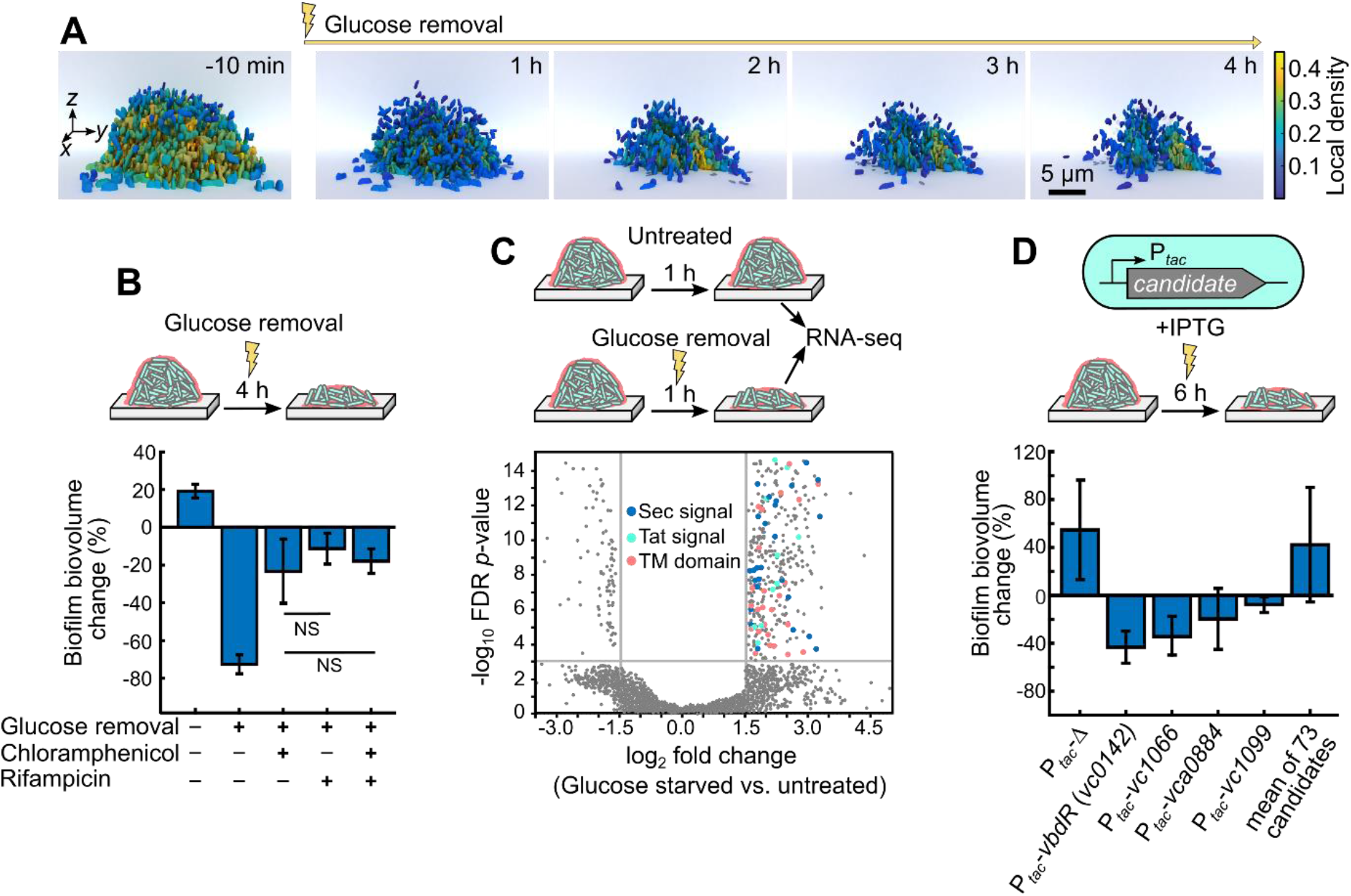
The biofilm dispersal regulator VbdR was identified in *V. cholerae* using transcriptomes and an over-expression screen. (A) *V. cholerae* biofilm dispersal dynamics at single-cell resolution during 4 h of carbon starvation. Each cell is colored according to the cell density in its local vicinity. Cell density was measured locally inside biofilms as the volume fraction of cell volume inside the biofilm volume, using BiofilmQ (61). Biofilms were grown in flow chambers and carbon starvation was achieved by exchanging the inflowing M9 medium to M9 without glucose. (B) Stressing pre-grown biofilms with glucose removal induces an active dispersal response that requires transcription and translation. Biofilm dispersal was quantified as the biofilm biovolume change during 4 h of glucose starvation with/without rifampicin (transcription inhibitor) or chloramphenicol (translation inhibitor). Chloramphenicol or rifampicin or both were added to biofilms 1 h prior to glucose removal. No significant difference (NS) was observed between conditions in which antibiotics were added. Error bars indicate SD of *n* ≥ 8 independent replicates for each condition. (C) Differentially regulated genes during biofilm dispersal were identified by comparing the transcriptome of biofilms that have undergone 1 h of glucose starvation and untreated control biofilms. Genes with absolute fold changes ≥ 3 and a false discovery rate (FDR) corrected *p*-value of ≤ 0.001 were considered to be differentially expressed. Upregulated genes were further screened for the presence of a secretory signal (Sec, blue dots), twin-arginine translocation signal (Tat, green dots), and transmembrane domains (TM, red dots). (D) All upregulated genes containing Sec, Tat, and/or TM were screened for their ability to induce biofilm dispersal, using plasmids with inducible expression constructs for each candidate based on P_*tac*_ and IPTG, which were introduced into *V. cholerae* strain KDV428 (WT with constitutive *sfgfp* expression). For each of the 77 candidates (see Table S1), the biofilm biovolume change during 6 h of IPTG induction was quantified and compared with the empty vector control. The four candidates that resulted in the strongest dispersal are shown here; data for the remaining 73 candidates are displayed in Fig. S1A. Error bars indicate SD of *n* ≥ 8 independent replicates for each condition.

For this glucose-removal-induced biofilm dispersal process, we sought to identify effectors that are responsible for disengaging the cells from the biofilm matrix. To discover potential effectors, we measured differential gene expression levels using RNA-seq for biofilm-bound cells that are in the process of actively dispersing, and a control treatment of non-dispersing biofilms (Fig. 1C). We hypothesized that dispersal effectors need to be either extracellular or possess a transmembrane domain, to be able to interact with the extracellular matrix. Of the 471 genes that are upregulated in the dispersing biofilms, only 77 genes code for proteins that contain either a secretion signal (Sec), twin-arginine translocation signal (Tat), or a transmembrane (TM) domain, which are listed in Table S1.

For each of these 77 candidates, we constructed a plasmid that uses the IPTG-inducible P_*tac*_ promoter to drive target gene expression, which was then moved into *V. cholerae*. To determine the ability of each of these candidates to cause dispersal, we induced the expression of each candidate in pre-grown biofilms for 6 h and quantified the resulting biofilm biovolume change (Fig. 1D, Fig. S1A). While the induction of many candidates caused no effect, or even increased biofilm growth (Fig. S1A), we identified 4 candidates whose induction caused dispersal (Fig. 1D): *vc0142*, *vc1066*, *vca0884*, and *vc1099*. Over-expression of these genes did not result in a substantial growth defect compared with an empty vector control (Fig. S1B). Strains deleted for these genes were still able to disperse with a similar capacity as the WT (Fig. S1C). Unexpectedly, none of these genes code for proteins with a known enzymatic domain (Fig. S2), yet all are highly conserved in *Vibrios*, except for *vc1066*, which is only conserved among *V. cholerae* strains. We speculate that these genes might code for regulatory proteins for which there is redundancy in the starvation-induced dispersal regulatory cascade. VC1099 possesses two TM domains, a conserved domain of unknown function (DUF412), and there are no characterized genes in its immediate neighborhood on the chromosome. VCA0884 also has not been characterized previously, but it contains a TM domain and its gene is located next to the *makDCBA* operon (motility-associated killing factor; *vca0880*-*vca0883*) (42). VC1066 is also uncharacterized but contains multiple TM domains and its gene is located in the immediate vicinity of *cdgH* (*vc1067*) coding for a diguanylate cyclase, which could influence c-di-GMP levels. Despite their proximity, *vc1066* and *cdgH* do not share one operon (43, 44). However, the strongest biofilm dispersal response was caused by the induction of *vc0142*, subsequently called *vbdR* for *Vibrio* biofilm dispersal regulator (Fig. 1D). VbdR is highly conserved among *Vibrio* species, and contains a conserved domain of unknown function and a transmembrane domain (Fig S2A). Given that *vbdR* induction is sufficient for causing biofilm dispersal and that it has the strongest effect on dispersal of all candidates, we focused on the mechanisms by which VbdR causes biofilm dispersal in this study.

Using a fluorescent protein transcriptional reporter and single-cell level imaging during biofilm dispersal, we confirmed that *vbdR* is induced during starvation-induced biofilm dispersal, and that it is not induced in control conditions (Fig. 2A), as expected from the transcriptome data (Fig. 1C). The spatiotemporal diagrams of *vbdR* expression also indicate that *vbdR* induction is stronger in the outer regions of the biofilm, where the cells are primarily dispersing (Fig. 1A). The fluorescent protein used for the reporter (mRuby3) has a slow maturation time (~150 min (45, 46)) so that the spatiotemporal measurements of *vbdR* expression indicate *vbdR* induction shortly after glucose removal. To explore the regulation of *vbdR*, we looked for binding motifs of known transcription factors in the *vbdR* promoter region and found binding motifs for HapR, a central transcription factor in quorum sensing, and CRP, involved in starvation response regulation (47, 48). By measuring a transcriptional reporter for *vbdR* in Δ*hapR* and Δ*crp* strains, we indeed observed reduced promoter activity compared to WT, indicating that *vbdR* is positively regulated by HapR and CRP (Fig. 2B). Published transcriptome data indicate that the *vqmR*/VqmA quorum sensing pathway also positively regulates *vbdR* (49). This regulation of *vbdR* is in line with our previous observation that quorum sensing and the starvation response jointly regulate *V. cholerae* biofilm dispersal (39).

**Figure 2.**
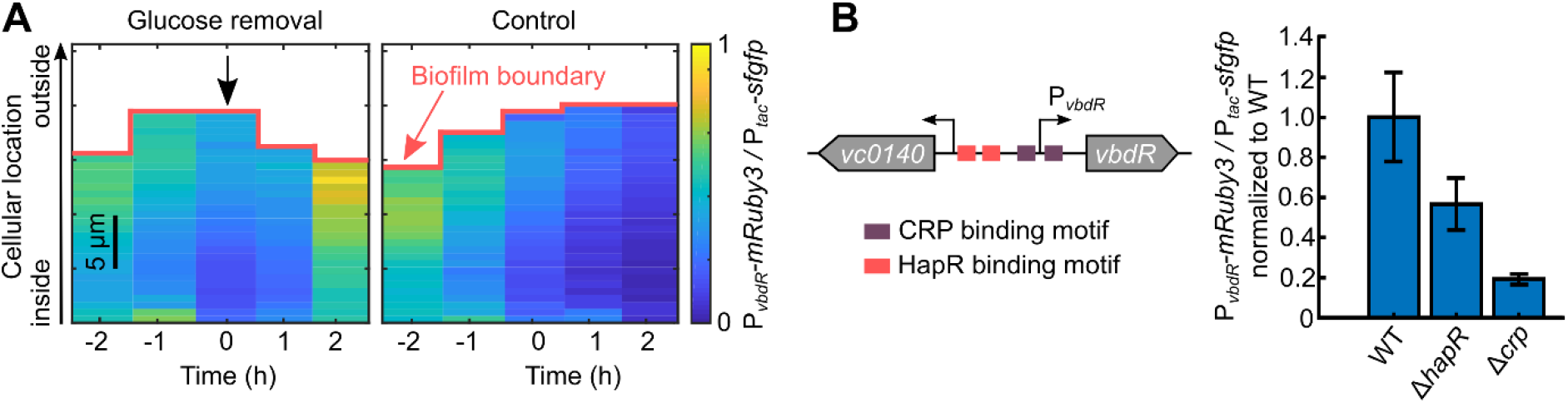
Characterization of *vbdR* expression and regulation. (A) The *vbdR* promoter reporter was measured in space and time inside biofilms, during carbon starvation-induced biofilm dispersal and during unperturbed biofilm growth. The kymograph *y*-axis represents the distance of each cell from the outer biofilm boundary facing the liquid medium; the *x*-axis indicates time, relative to the glucose removal event (indicated by the black arrow at *t* = 0 h). The *vbdR* promoter reporter was quantified for each cell in the biofilm, by normalizing the fluorescence of mRuby3 (expressed from the *vbdR* promoter) by the fluorescence of sfGFP (expressed from the constitutive P_*tac*_ promoter). The maturation time of mRuby3 is ~150 min (45, 46). Each heatmap is representative of *n* = 10 independent replicates. (B) Putative binding sites for HapR and CRP on the *vbdR* promoter are indicated. The *vbdR* promoter reporter (P_*vbdR*_-*mRuby3* signal divided by the P_*tac*_-*sfgfp* signal, normalized to the WT) was measured in the wild type, Δ*hapR*, and Δ*crp* strains, for cells grown in liquid shaking culture. These results indicate that the major transcription factors HapR and CRP regulate *vbdR* transcription. Error bars indicate SD of *n* = 3 independent replicates, each with hundreds of cells.

To study how VbdR causes cellular dispersal from biofilms, we first quantified the dispersal dynamics following over-expression of *vbdR*. In pre-grown biofilms (diameter of 20-25 μm), induction of *vbdR* expression caused reduced cell density at all spatial locations in the biofilm, and simultaneously caused a shrinkage of the outer biofilm boundary (Fig. 3A). We previously observed that a cell density reduction in biofilms can be mediated by a modification of the extracellular matrix (37), which led us to investigate whether the matrix undergoes significant changes as a result of *vbdR* production. Using antibodies or lectins conjugated to fluorescent dyes, we therefore quantified the abundance of the major known matrix components RbmA, RbmC, Bap1, and VPS, in a 0.6 μm-thick shell around each cell in the biofilm in space and time during *vbdR* over-expression (Fig. 3B). These results show that the levels of RbmC and Bap1 were unchanged in the shell around the cells. In contrast, these data also show that the RbmA and VPS abundances in the shell around the cells decreased during induced over-expression of *vbdR*, compared to an empty vector control (Fig. 3B). VPS accounts for ~50% of the extracellular matrix in biofilms (21, 50), yet the combined staining with fluorescently labeled Concanavalin A and wheat germ agglutinin lectins yields a much weaker fluorescence signal than the immunofluorescence signal of the matrix proteins, which may be due to an incomplete labeling of VPS. We therefore expect that the matrix protein immunofluorescence data is a more accurate reflection of the true matrix dynamics than the lectin staining.

**Figure 3.**
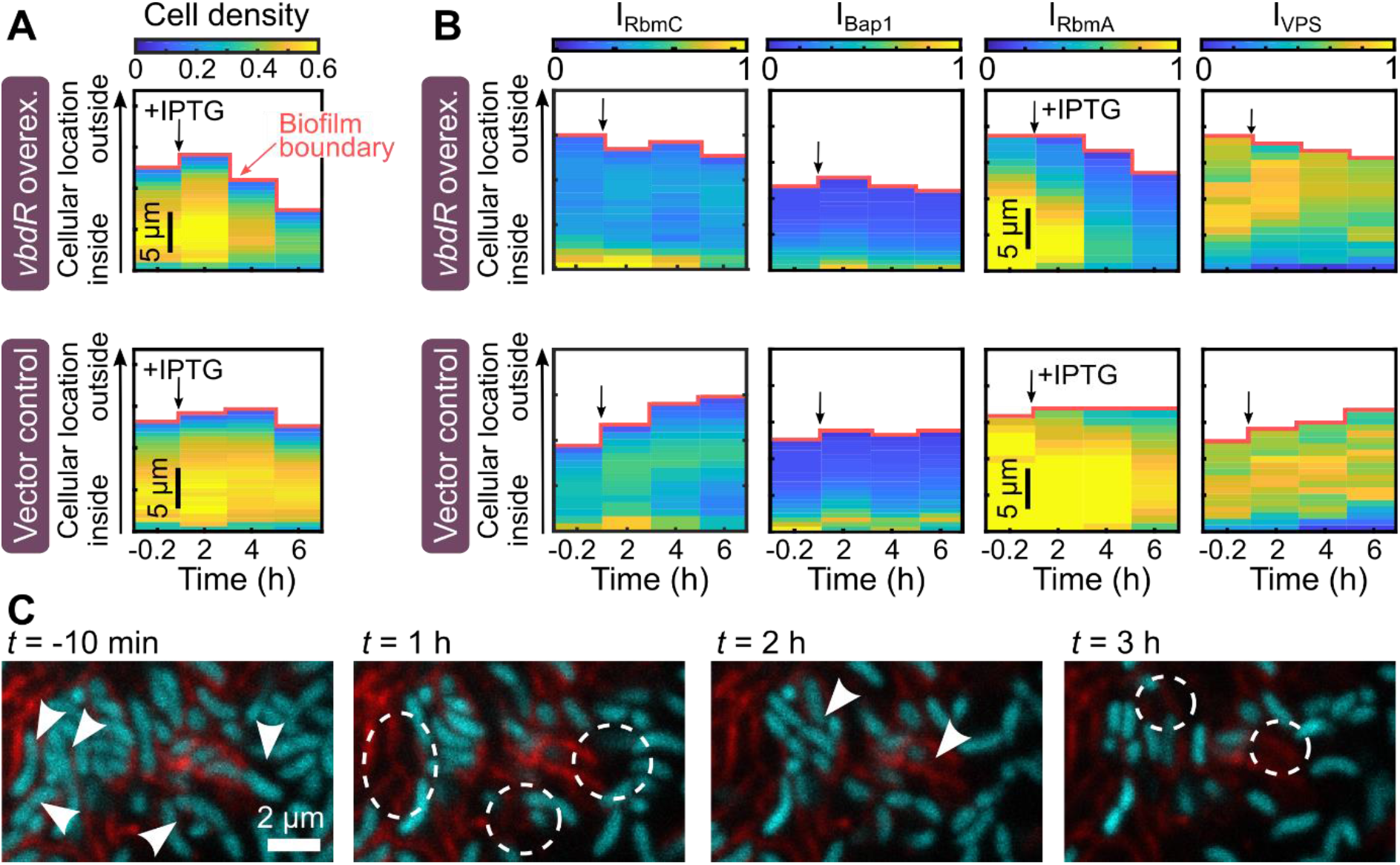
*vbdR* over-expression reduces biofilm cell density and modifies matrix composition. (A) Spatiotemporal measurements of the biofilm-internal cell density during *vbdR* over-expression from plasmid pNUT1594 using the IPTG-inducible P_*tac*_ promoter (top heatmap) or an empty vector control pNUT1246 (bottom heatmap). Cell density was measured locally inside biofilms as the volume fraction of cell volume inside the biofilm volume, using BiofilmQ (61). After growing biofilm colonies up to a diameter of 20-25 μm in M9 medium, *vbdR* over-expression was achieved by exchanging the feeding tubing and syringes to contain M9 + 1 mM IPTG immediately after imaging the first time point. The kymograph *y*-axis corresponds to the distance of each cell from the biofilm outer boundary; measurements from cells with similar distances to the biofilm boundary are averaged. A reduction of the biofilm outer boundary during *vbdR* over-expression reflects dispersal. Each kymograph is representative of *n* = 10 independent replicates. (B) Spatiotemporal measurements of biofilm matrix component abundance during *vbdR* over-expression (top row) and an empty vector control (bottom row), using immunofluorescence for the matrix proteins RbmC, Bap1, and RbmA, and FITC-conjugated wheat germ agglutinin and Concanavalin A lectins for VPS. The heatmaps show I_matrix_ for the individual matrix components, which was measured by quantifying the immunofluorescence or lectin fluorescence in a shell (0.6 μm thickness) around each cell, for all cells in the biofilm. The I_matrix_ signal is scaled in linear arbitrary units between 0 and 1. For matrix protein immunofluorescence, a 6xHis tag was added to the native chromosomal locus of *rbmA* and *rbmC*, and a HA tag was added to *bap1*. Anti-His or anti-HA antibodies conjugated to Alexa Fluor dyes were used to label proteins in living biofilms (20). Each kymograph is representative of *n* = 10 independent replicates. (C) Confocal microscope image time series of cells (cyan) leaving from pockets of RbmA (red immunofluorescence) in biofilms during dispersal. Arrow heads point to cells that are leaving in the next image frame; dashed lines indicate regions where cells have left. Cells that have dispersed leave behind empty shells of RbmA.

The reduction of RbmA and VPS abundance in the shell surrounding each cell could be due to RbmA and VPS degradation, or due to an increased spatial separation between RbmA and VPS and the cells, or due to both effects. Interestingly, high-resolution confocal microscopy images indicate that after cells disperse from the biofilm, they leave behind empty shells of RbmA-containing matrix (Fig. 3C). In addition, these images showed that the overall RbmA immunofluorescence decreased during *vbdR* over-expression (Fig. 3C). Together, the results from Fig. 3B, C indicate that the cells separate from the RbmA matrix shell, and that RbmA levels decrease during cellular departure.

Bioinformatic analysis of VbdR did not reveal a catalytic domain that could directly process RbmA or VPS. Our transcriptome data also indicated that VbdR does not change the transcription of the *vps*-I and *vps*-II operons or *rbmA*, which we confirmed using a *rbmA* transcriptional reporter (Fig. S3). We therefore hypothesized that VbdR functions as a regulator that activates downstream effectors which cause the separation between the cells and their matrix shell and matrix degradation. To determine the VbdR regulon in *V. cholerae*, we performed differential gene expression analysis based on RNA-seq measurements from biofilms that have undergone 1 h of *vbdR* induction, compared with a control treatment (Fig. 4A). This resulted in only 8 genes that were significantly upregulated. The highest fold change was displayed by *pilT* and *pilU*, which code for type IV pilus retraction ATPases (28, 51–53). Interestingly, strains deleted for 7 of the 8 effector candidates showed no change in their ability to display starvation-induced biofilm dispersal, compared with the WT (Fig. 4B). A Δ*pilT* strain could not be included in Fig. 4B because cells lacking *pilT* did not form biofilms in flow chambers, which has previously been observed in other *V. cholerae* strain backgrounds (54). To determine the effect of PilT abundance on biofilm dispersal, we generated an inducible *pilT* knock-down strain based on CRISPRi, using a single guide-RNA targeting the *pilT* promoter. These experiments showed that *pilT* knock-down after the initial surface-attachment of cells caused a strong defect in the dispersal ability, compared with uninduced conditions or the empty vector control (Fig. 4C). Therefore, out of the 8 VbdR-upregulated proteins, only decreased PilT levels resulted in a dispersal defect.

**Figure 4.**
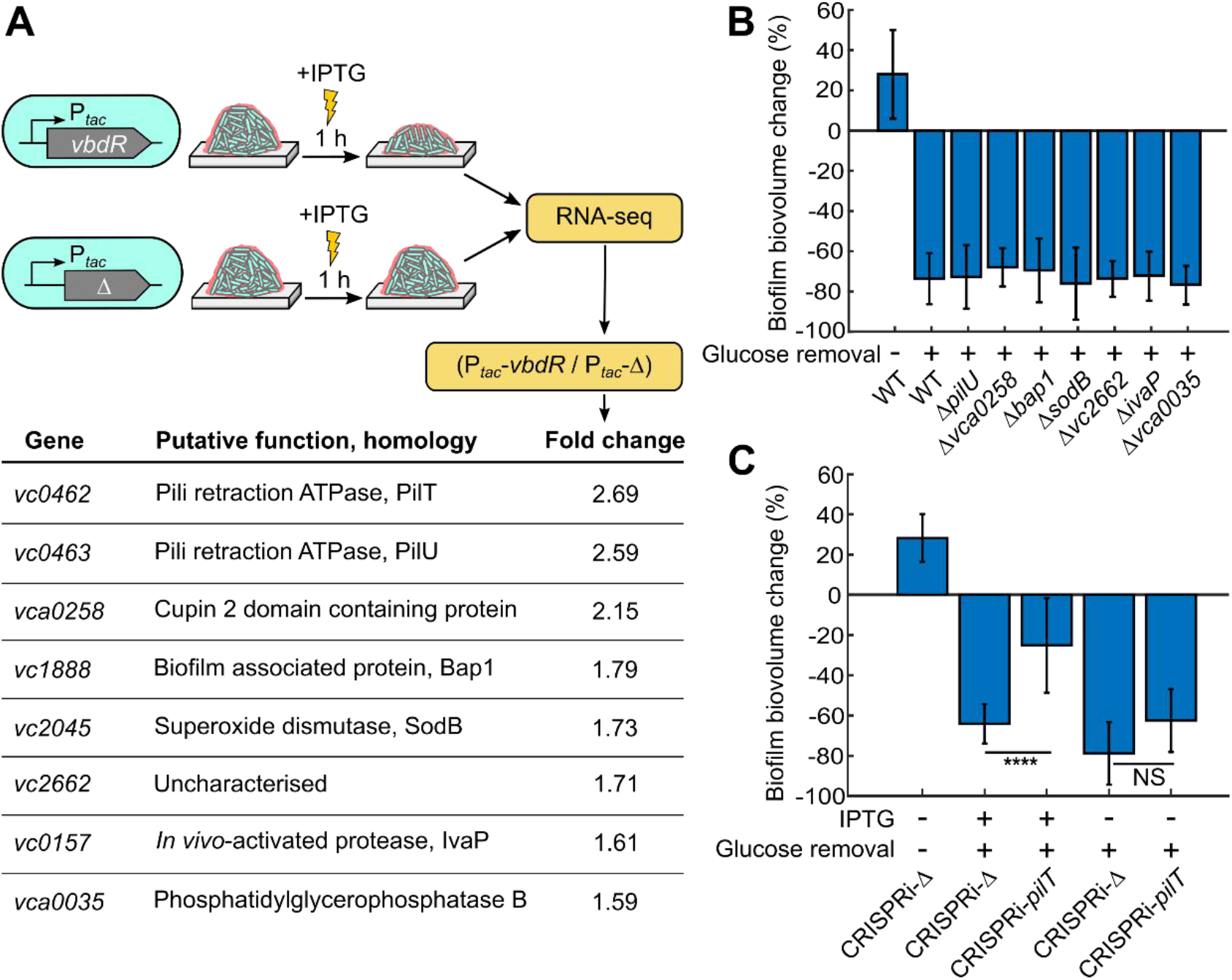
Identification of genes that are upregulated by VbdR and their effect on biofilm dispersal. (A) To determine the regulon of VbdR, biofilms of a strain that harbored an IPTG-inducible *vbdR* construct, or a strain that harbored an empty vector control were grown for 20 h in flow chambers. After 1 h of IPTG induction, biofilms were harvested and their transcriptome was determined using RNA-seq. Upregulated genes with an FDR corrected p-value of ≤ 0.001 are listed by descending fold change. (B) Using the glucose removal assay, the biofilm dispersal capability was determined for strains carrying a single deletion of the genes listed in panel A, except for *pilT* (*vc0462*) – Δ*pilT* strains do not form biofilms. As control experiments, the glucose removal assay was also performed for the WT, and the biofilm biovolume change was measured for the WT without removing glucose. Error bars indicate SD of 30 biofilms in *n* = 3 independent replicates for each condition. (C) Using the glucose removal assay, the biofilm dispersal capability was determined for *pilT* knock-down strains. An IPTG-inducible CRISPRi-*pilT* construct with a guide RNA targeting the *pilT* promoter region was used to knock down *pilT* transcription, and results are compared to an empty vector control CRISPRi-Δ. In these experiments, IPTG was added to the media 6 h after initial attachment of cells on the glass surface, and biofilms were grown until they reached a diameter of 20-25 μm before removing glucose to trigger dispersal. The biofilm biovolume change was measured before glucose removal and 4 h into the glucose removal treatment. As a control, the biovolume change was also measured without removing glucose. Error bars indicate SD of 30 biofilms in *n* = 3 independent replicates for each condition; **** indicates *p*<0.0001.

How does PilT contribute to active biofilm dispersal? Characterizations of the type IV pilus retraction ATPases PilT and PilU have recently revealed that PilT is the primary ATPase, yet PilU can contribute in a PilT-dependent manner in conditions when a high retraction force is required (28, 51, 52, 55). In *V. cholerae*, PilT and PilU are responsible for the retraction of the DNA-uptake/competence pilus (formerly called chitin-regulated pilus) and the MSHA pilus. The third type IV pilus in *V. cholerae*, the toxin-coregulated pilus, relies on the minor pilin TcpB for retraction without an ATPase (56). Which type IV pilus does PilT control in biofilms? Our transcriptome data from WT *V. cholerae* (used for Fig. 1C) showed that out of the three type IV pilus systems, the MSHA pilus has by far the most transcripts inside biofilms (Fig. 5A). To test if MSHA pili are indeed present inside biofilms, we used immunofluorescence based on a fluorescent anti-His antibody and a translational fusion of the major pilin MshA to a 6xHis tag. Confocal microscopy images revealed that MSHA pili are highly abundant at all locations inside the biofilm (Fig. 5B). Using fluorescently labeled maleimide binding to MshA, into which an exposed cysteine residue has been introduced (28, 51, 57), we performed experiments that confirmed this finding (Fig. S4). Matrix secretome measurements have previously hinted that MshA is present inside biofilms (25), and it was shown that MSHA pili are important for initial surface attachment and detachment of individual *V. cholerae* cells (28, 30–32, 54, 58). In line with these reports, our findings now directly demonstrate that MSHA pili are an abundant extracellular matrix component of 3D *V. cholerae* biofilm colonies. Given the abundance of MSHA pili in biofilms, the upregulation of *pilT* by VbdR during biofilm dispersal, and the requirement of PilT for full dispersal (Fig. 4C), we hypothesized that the retraction of MSHA pili is necessary during biofilm dispersal.

**Figure 5.**
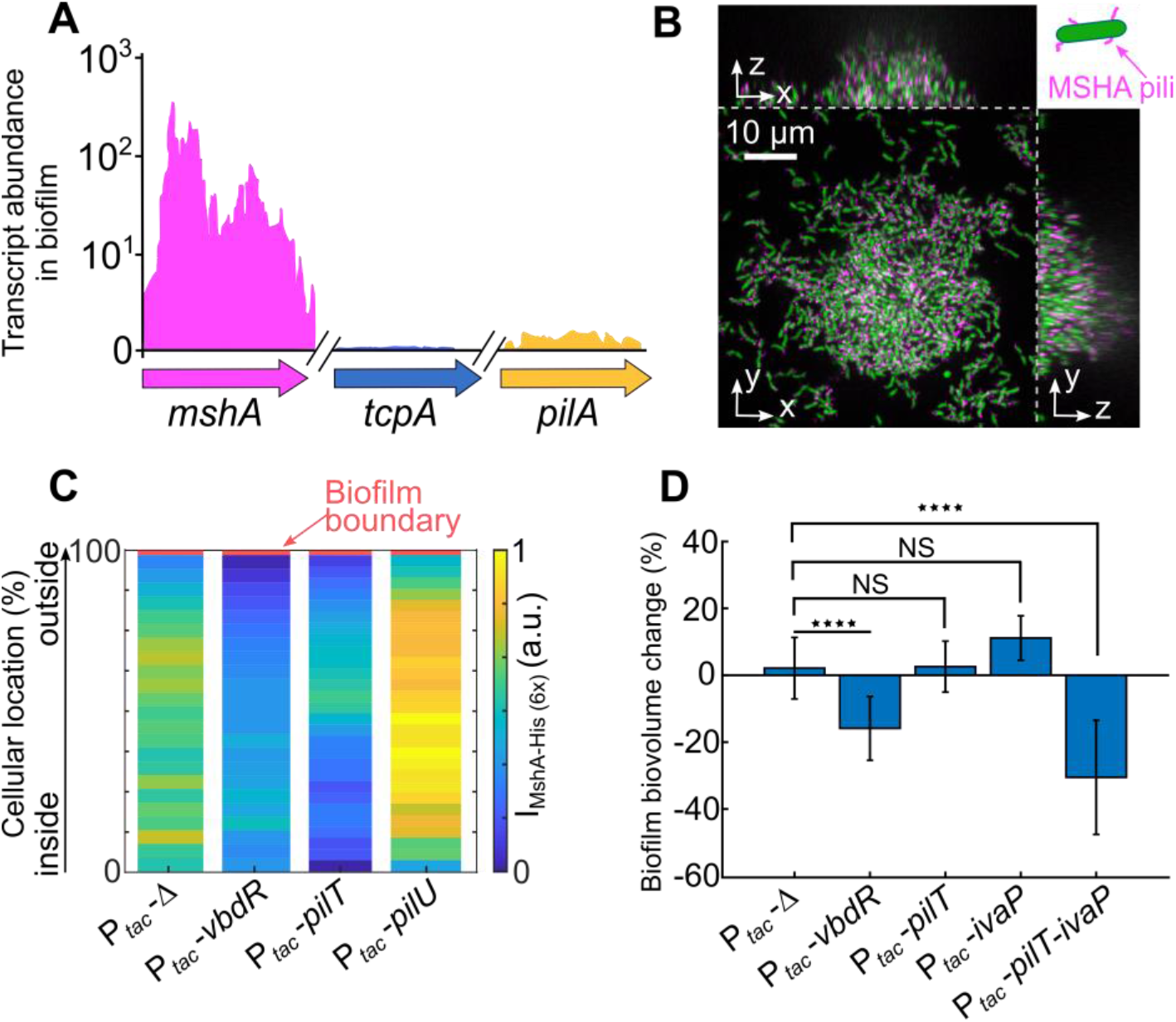
The joint effect of MSHA pilus retraction and IvaP-based matrix degradation is sufficient for causing biofilm dispersal. (A) RNA-seq measurements of biofilm colonies were used to compare the transcript abundance of the major pilins of all type IV pili in *V. cholerae*. (B) Immunofluorescence shows that MSHA pili (magenta) are present in live biofilms; cells constitutively express sfGFP (green, strain KDV2451). To perform immunofluorescence, the native *mshA* gene was replaced by a *mshA-6xHis* translational fusion, and anti-His antibody conjugated to Alexa Fluor 555 was added to the medium during the entire growth period. A schematic diagram illustrates the fluorescence colors of a cell and MSHA pili. (C) Quantification of MSHA pili abundance in biofilms after over-expression of *vbdR*, *pilT*, or *pilU*. The MSHA pilus abundance was quantified by measuring the antibody fluorescence in a shell around each cell (using the BiofilmQ software), and pilus abundance measurements were performed with spatial resolution in the biofilm, as a function of distance from the outer biofilm boundary with the liquid growth medium. For these experiments, biofilms were grown for 18 h before *vbdR*, *pilT*, *pilU*, or the empty vector control were induced with IPTG for 5 h. After 4 h of IPTG-induction, fluorescent anti-His antibodies were added to label MSHA-His pili to allow the antibodies to diffuse into the biofilm for 1 h, followed by 3D confocal imaging. Each spatial abundance map of MSHA pili is the average of 20 biofilms from *n* = 3 independent replicates. P_*tac*_-Δ represents the empty vector control. (D) Biofilm dispersal was quantified during 6 h of IPTG induction for strains harboring different plasmids with IPTG-inducible expression constructs. Glucose was always kept in the growth medium. The joint expression of *pilT* and *ivaP* induced biofilm dispersal even in the presence of glucose. Error bars indicate SD of *n* = 10 replicates for each condition; **** indicates *p*<0.0001.

To test whether *vbdR* expression actually causes MSHA pilus retraction in biofilms, we used confocal microscopy and computational image analysis to quantify the MSHA pili abundance in a shell around each cell of the biofilm, for biofilms in which we ectopically induced expression of *vbdR*, *pilT*, *pilU*, or an empty vector control for 6 h (Fig. 5C). Induction of both *vbdR* and *pilT*, but not *pilU*, resulted in significantly reduced MSHA pili levels in biofilms, compared to the control (Fig. 5C). Given that MSHA pilus retraction cannot degrade the other matrix components, the requirement of MSHA pilus retraction for full biofilm dispersal indicates that MSHA pilus retraction releases the cells from the existing matrix during dispersal.

For MSHA pilus retraction to cause a release of cells from the matrix, the MSHA pilus needs to bind to at least one component of the matrix or other cells in the biofilm prior to pilus retraction. We therefore investigated if the major pilin MshA has protein binding partners in the biofilm matrix. To this end, we fused a GST tag to MshA, and performed a pull-down of purified MshA-GST mixed with purified biofilm matrix, followed by a mass spectrometry-based proteomics (see Fig. S5 for workflow). As a control, we used an identical workflow but with purified GST protein instead of MshA-GST as bait. The proteomics analysis identified 29 proteins enriched in MshA-GST experiments (Table S2). From these 29 proteins, 22 proteins are predicted to be localized in the cytoplasm, which indicates the presence of lysed cells in our purified matrix. Lysed cells naturally occur during *V. cholerae* biofilm growth (23, 37, 59), but they could also arise from our matrix purification protocol (see Materials and Methods). As MSHA pili have been shown to bind to many abiotic and biotic surfaces non-specifically (27, 31, 58, 60), these results are consistent with the idea that MSHA pili bind to several proteins non-specifically. It is also known that MSHA pili bind to the polysaccharides mannose and chitin (27, 31), and we speculate that MSHA pili could potentially bind to VPS in the matrix (60). These findings suggest that MSHA pili have numerous non-specific binding partners in biofilms.

Given that the *pilT*-mediated MSHA-pilus retraction is required for full biofilm dispersal, we tested if this pilus retraction process is also sufficient for biofilm dispersal, by measuring biofilm dispersal following *pilT* over-expression. However, *pilT* over-expression did not cause biofilm dispersal (Fig. 5D). Therefore, another process must simultaneously occur to enable dispersal. Figure 3C showed that the RbmA abundance decreases during VbdR-induced dispersal, and Fig. 4A showed that *vbdR* over-expression causes an upregulation of *ivaP* (Fig. 4A), which codes for a secreted serine protease that is known to process RbmA and other proteins (33, 34). We therefore tested if *ivaP* induced over-expression causes biofilm dispersal, yet these results were negative (Fig. 5D). However, inducing the expression of both *pilT* and *ivaP* together with IPTG did result in biofilm dispersal (Fig. 5D). None of these over-expressions caused a substantial growth rate defect, compared with an empty vector control (Fig. S6). The joint effect of MSHA pilus retraction and RbmA degradation *via* IvaP is therefore sufficient for causing biofilm dispersal.

## Discussion

Our results indicate that during *V. cholerae* biofilm dispersal, MSHA pilus retraction releases cells from the potentially numerous binding partners in the matrix. Simultaneously, RbmA degradation opens up the matrix shell around the cells to enable cellular departure from the biofilm following MSHA pilus retraction. Given that IvaP is not required for dispersal (Fig. 4B), other enzymes need to also be able to degrade the matrix shell around the cells. Irrespective of how the matrix shell is opened up, the release of cells from the matrix *via* MSHA-pilus retraction is a key process that is necessary for full dispersal (Fig. 4C).

More generally, the commitment of cells to form biofilms is an energetically costly investment for the cells as it involves the production of copious amounts of extracellular matrix, and results in a lower single-cell growth rate of the biofilm-bound cells compared with planktonic cells (9, 10). It is therefore not surprising that biofilm formation is a highly regulated process on all levels, involving external signals, second messenger signals, two-component systems, regulatory RNA, as well as transcriptional and post-transcriptional regulation (21). Given this complex regulation of biofilm formation, it is likely that biofilm dispersal is a phenotype with similarly complex regulation. Our previous results have shown that *V. cholerae* biofilm dispersal in flow chambers requires the integration of both nutrient starvation and quorum sensing regulation, to ensure robust cellular decisions about dispersal (39). Here we have shown that the regulation of biofilm dispersal is more complex, finding several additional putative regulators whose induction is sufficient to mediate dispersal (Fig. 1D). In this study, we identified and characterized VbdR and demonstrated that it causes dispersal *via* MSHA pilus retraction and matrix degradation. However, investigations of the other putative regulators VC1066, VCA0884, and VC1099, as well as the role of c-di-GMP will likely yield additional levels of regulation and additional effectors. A characterization of the regulators and effectors involved in different biofilm dispersal model systems will not only lead to an improved understanding of dispersal, but also to further insights into the functionality and properties of the biofilm matrix.

## Conclusion

Our work has leveraged the increasing knowledge about *V. cholerae* matrix composition and function to demonstrate that the retraction of MSHA pili in combination with matrix degradation results in a release of the cells from the matrix, which underlies the departure of cells from biofilms during biofilm dispersal. This important role of MSHA pili during biofilm dispersal stems from the high abundance of MSHA pili in the matrix, which anchor cells in the matrix during biofilm growth. We discovered that MSHA pilus retraction is triggered by the upregulation of VbdR, a novel biofilm dispersal regulator that is controlled by quorum sensing and CRP. Together, these results provide a detailed mechanistic understanding of the *V. cholerae* biofilm dispersal process during carbon starvation. Biofilms are notoriously difficult to prevent or eliminate, yet approaches based on manipulating the endogenous dispersal mechanisms of biofilms could be technologically very useful for controlling biofilms.

## Acknowledgements

We are grateful to Alexander Lepak and Gert Bange for help with protein purification, Lucia Vidakovic, Keerthana Raveendran, and Sanika Vaidya for strain construction and important discussions, Kyle Floyd and Fitnat Yildiz for advice on MSHA pili, Raimo Hartmann for contributions to image analysis at the early stage of the project, and Witold Szymanski for advice regarding proteomic mass spectrometry. This work was supported by grants from the Alexander von Humboldt Foundation (to Ka.No.), Studienstiftung des deutschen Volkes (H.J.), Deutsche Forschungsgemeinschaft (SFB987, DR 982/5-1, and EXC 2051 - Project‐ID 390713860), Minna James Heineman Foundation, European Research Council (StG-716734 and StG-758212), Bundesministerium für Bildung und Forschung (TARGET-Biofilms), Max Planck Society, and the Swiss National Science Foundation NCCR “AntiResist” (to K.D.).

## Materials and Methods

### Media, strains, culture conditions

All *V. cholerae* strains used in this study are derivatives of the wild type *V. cholerae* O1 biovar El Tor strain C6706, which is capable of quorum sensing (62). All *V. cholerae* strains were grown in liquid LB medium supplemented with appropriate antibiotics (gentamycin 30 μg/mL, kanamycin 100 μg/mL) at 28 °C for routine growth. Biofilm experiments with *V. cholerae* were performed in M9 minimal medium, supplemented with 2 mM MgSO_4_, 100 μM CaCl_2_, MEM vitamins, 0.5% glucose, and 15 mM triethanolamine (pH 7.1). If necessary, 1 mM IPTG was added to the media to induce the P_*tac*_ promoter, respectively. Detailed lists of strains and plasmids are provided in Table S3 and Table S4, respectively.

### Cloning methods

To construct plasmids and bacterial strains, standard molecular biology techniques were applied (63). All enzymes for cloning were purchased from New England Biolabs or Takara Bio. *V. cholerae* deletion mutations were genetically engineered using derivatives of the pKAS32 suicide vector harbored in *E. coli* S17-λ pir (64). All over-expression constructs were cloned into a low copy number plasmid with a pSC101* origin of replication and a gentamycin resistance cassette, or inserted at the chromosomal *lacZ* site with the help of the suicide plasmid pKAS32. Plasmid clones were first constructed in *E. coli* Top10 and then mated into *V. cholerae* using an *E. coli* strain harboring the conjugation plasmid pRK600. Detailed cloning strategies are described below. Oligos are listed in Table S5 and were commercially synthesized by Eurofins or Sigma Aldrich. All plasmid constructs that were created were sequenced by Eurofins for their correctness.

### Plasmids for inducible over-expression

For constructing inducible over-expression plasmids for putative dispersal-inducing factors, we used a standard plasmid backbone, pNUT1246. The plasmid pNUT1246 contains a gentamicin resistance cassette, the pSC101* origin of replication, the IPTG-inducible promoter P_*tac*_, and the *lacI*^Q1^ repressor to tightly repress the expression from P_*tac*_ in the absence of IPTG. Gibson assembly was used to generate the final constructs for all plasmids (65). To simplify the cloning of the over-expression plasmids, the oligos kdo1898/kdo1899 were used to amplify the pNUT1246 backbone, and optimal overlapping nucleotide bases were defined for all other oligos. These plasmids were then introduced into strain KDV428. A list of the plasmids and primers with detailed information about the cloned genes is provided in Tables S4 and Table S5.

### Plasmids for chromosomal gene deletions

To generate gene deletions in *V. cholerae,* plasmids based on the suicide vector pKAS32 were generated for respective genes. Briefly, the vector pNUT144 (a derivative of pKAS32) was amplified using oligos kdo1968/kdo1969, and 1 kb upstream and 1 kb downstream of the gene of interest were amplified with primers specific to the gene locus of interest (listed in Table S5). The vector backbone and the inserts were combined using the standard Gibson assembly protocol (65)

### Reporter strains

To generate the plasmid pNUT2172, which is a *mRuby3*-based transcriptional reporter for *vbdR*, the promoter sequence of *vbdR* was amplified with oligonucleotides kdo2741/kdo2981 and cloned upstream of *mRuby3* in the plasmid pNUT1029. The reporter plasmid pNUT2172 was conjugated into strains KDV428, KDV433, and KDV2624 to generate strains KDV2350, KDV2365, and KDV2625, respectively.

### Flow chamber biofilm experiments

*V. cholerae* biofilms were grown in microfluidic flow chambers made from polydimethylsiloxane (PDMS) and glass coverslips, as described earlier (39). PDMS and glass coverslips were bonded using an oxygen plasma, and resulted in flow chambers with dimensions 7000 μm length, 500 μm width, 100 μm height. The microfluidic design contained either 4 or 8 channels of identical dimensions, which are independent from each other. The manufacturing process of these microfluidic channels guarantees highly reproducible channel dimensions and surface properties in the channels. The channels were imaged on an inverted microscope, through the coverslip at the bottom of the channels. Each channel was inoculated with freshly-diluted cultures from *V. cholerae* strains, which were grown overnight at 28 °C in liquid LB medium under shaking conditions. Following inoculation of the channels, the cells were given 1 h to attach to the glass surface of the channel without flow, before a flow of 100 μL/min M9 medium was initiated for 45 seconds to wash away non-adherent cells and to remove LB medium from the channels. The flow rate was then set to 0.1 μL/min until the end of the experiment, and the flow chambers were incubated in a 25 °C incubator. Flow rates were controlled using a high-precision syringe pump (Harvard Apparatus).

### Flow chamber biofilm dispersal assay

To induce biofilm dispersal, we used one of two assays. In one assay, we used carbon starvation to trigger dispersal. For this, we grew biofilms up to a well-defined size (20-25 μm colony diameter) in the flow chambers at 25 °C, followed by the exchange of the inflowing M9 medium to M9 medium without glucose. This exchange of the medium was achieved by exchanging the syringe and tubing that feeds medium into the microfluidic channel. A 3D confocal image of each biofilm was acquired immediately before the exchange of the syringes and 4 h after the exchange of the syringe, to quantify the dispersed biofilm biovolume using the BiofilmQ software tool for image analysis (61).

In the other assay, we tested if the over-expression of a particular gene could induce biofilm dispersal. For this, biofilms were grown up to a well-defined size (20-25 μm colony diameter) in flow chambers at 25 °C, followed by the exchange of the inflowing M9 medium to M9 medium with 1 mM IPTG. In a separate flow channel on the same microfluidic chip, biofilms of a *V. cholerae* control strain (KDV911, which is the KDV428 strain containing the empty vector pNUT1246) were exposed to the same treatment, as a control in every experiment. Confocal 3D images of biofilms were taken immediately before the exchange of syringes. After induction with IPTG for 6 h, dispersed or loosely attached cells were washed away with 20 μL/min flow for 10 min before taking the final confocal 3D image. A comparison of the initial and final 3D images was used to quantify the biofilm biovolume change (as a measure for biofilm dispersal), using the BiofilmQ software.

### *vbdR* reporter imaging in biofilms and in liquid shaking culture

Biofilms of strain KDV2350 (*lacZ*::P_*tac*_-*sfgfp,* pNUT2172) were grown at 25 °C under constant flow (0.1 μL/min) in flow channels, as described above, and we define time *t* = 0 as the time point when the flow was set to 0.1 μL/min. At *t* = 20 h biofilms were imaged in 3D with a confocal microscope, followed by another 3D image at *t* = 21 h. After the second imaging round, biofilms were induced for dispersal by replacing the inflowing M9 medium by M9 without glucose. Imaging was continued for another 2 h with 1 h intervals.

Liquid shaking cultures of KDV2350 (genotype: *lacZ*::P_*tac*_-*sfgfp,* pNUT2172), KDV2365 (genotype: *lacZ*::P_*tac*_-*sfgfp, ΔhapR,* pNUT2172), and KDV2625 (genotype: *lacZ*::P_*tac*_-*sfgfp, Δcrp,* pNUT2172) were grown in LB at 28 °C under shaking conditions until OD_600_ = 2. From this culture, 10 μL was spotted on a glass coverslip and a thin slice of LB agar was added on top of the culture to inhibit movement of cells for imaging with confocal microscopy.

### Sample collection for transcriptomes of dispersed biofilms

To collect a sufficient amount of biomass for RNA-seq, we grew *V. cholerae* C6706 (KDV428) biofilms in our microfluidic flow chambers at 25 °C in six separate, identical channels. The biomass collection was performed as follows: After growth for 21 h, the inflowing M9 medium with 0.5% glucose was exchanged to M9 without glucose for three of the six channels (starved biofilms), by exchanging the tubing and syringes at the channel inlet. For the other three channels, the tubing and syringes were also exchanged but the inflowing medium was kept as M9 medium with glucose (untreated control biofilms). After 1h, a mixture of 47.5% (vol/vol) EtOH, 2.5% (vol/vol) phenol, and 50% M9 medium(vol/vol) was flown through the flow chamber channels to terminate transcription and translation. Then, the PDMS was removed from the coverslip and the biofilms were scraped off the coverslip using a clean razor blade. Biofilms from all six channels were collected in separate collection tubes, which were snap-frozen in liquid nitrogen and samples were stored at −80 °C until RNA isolation. This process was repeated on two separate days, to obtain a total of six samples for each of the two conditions.

### Sample collection for transcriptomes of biofilm cells after *vbdR* induction

Biofilms of *V. cholerae* C6706 strains KDV911 and KDV1112, which harbor either the empty vector (pNUT1246) or the *vdbR* over-expression plasmid (pNUT1594), respectively, were grown under constant flow of M9 medium supplemented with 0.5% glucose at 25 °C in six separate, identical channels. Three channels contained biofilms of KDV911, and three channels contained biofilms of KDV1112. After unperturbed biofilm incubation for 20 h, the expression of the *vdbR* gene was induced by replacing the inflowing M9 medium with M9 containing 1 mM IPTG for all six channels. After induction for 1 h, biofilm cells were harvested adding a mixture of EtOH and phenol to a final concentration of 47.5% (vol/vol) EtOH, 2.5% (vol/vol) phenol, and 50% (vol/vol) M9 medium, followed by PDMS removal and scraping biofilms off the glass coverslip and snap‐freezing cells in liquid nitrogen. Samples were stored at −80 °C until RNA isolation. This process was repeated on three days, to obtain a total of nine samples for each of the two strains.

### RNA-seq and transcriptome analysis

Total RNA was isolated using the hot phenol method, as described previously (66) and DNA was digested with TURBO DNase (Thermo Fischer Scientific). Ribosomal RNA was depleted using the Ribo-Zero kit for Gram-negative bacteria (Illumina) and RNA integrity was confirmed using automated electrophoresis (BioAnalyzer, Agilent). Directional cDNA libraries were prepared using the NEBNext Ultra II Directional RNA Library Prep Kit for Illumina (E7760, NEB). The libraries were sequenced using a HiSeq 1500 or 3000 system in single-read mode for 100 or 150 cycles. The read files were imported into CLC Genomics Workbench v10.1.1 (Qiagen) and trimmed for quality and 3’ adaptors. Reads were mapped to the *V. cholerae* reference genome (NCBI accession numbers: NC_002505.1 and NC_002506.1) using the “RNA-Seq Analysis” function in the CLC software with standard parameters. Reads mapping to annotated coding sequences were counted, normalized (counts per million, CPM) and transformed (log_2_). Differential expression between the conditions was tested using the “Differential Expression for RNA-seq” command in the CLC software. Genes with a read count < 10 in any condition were excluded from analysis. Genes with a fold change ≥ 3.0 and a false discovery rate (FDR) adjusted *p*-value ≤ 0.001 were defined as differentially expressed. For the analysis of RNA-seq results from the *vbdR* pulse-induction, genes with a fold change ≥ 1.5 and a FDR adjusted p-value ≤ 0.001 were defined as differentially expressed.

### *In-silico* analysis of potential dispersal effectors

All 473 upregulated genes from RNA-seq data obtained from biofilm samples with/without glucose starvation were screened for the presence of either a Sec signal sequence, twin-arginine translocation (Tat) signal sequence, or transmembrane domain (TM) with the help of online available bioinformatics tools SignaIP (67), TatP (68), and TMM (69), respectively. The resulting list of all 77 upregulated genes with Sec, Tat, or TM sequences is provided in Table S1.

### *vbdR* promoter characterization

To identify putative promoter for *vbdR*, we used the online software tool BROM (70). The CRP binding site on the *vbdR* promoter was identified with the help of published literature (71) HapR binding sites on the *vdbR* promoter were identified manually by aligning the consensus binding sequence for HapR (72). The CRP and HapR binding sites are show schematically in Fig 2.

### VbdR sequence alignment

VbdR (vc0142) amino acid sequences from various *Vibrio* species were aligned using the ClustalW alignment webtool (https://www.ebi.ac.uk/Tools/msa/clustalo/), using the following sequences: *Vibrio cholerae* (NCBI txid948564), *Vibrio mimicus* (NCBI txid1267896), *Vibrio anguillarum* (NCBI txid55601), *Vibrio vulnificus* (NCBI txid672), *Vibrio alginolyticus* (NCBI txid663), *Vibrio parahaemolyticus* (NCBI txid670), *Vibrio metoecus* (NCBI txid1481663), *Vibrio mexicanus* (NCBI txid1004326), *Vibrio cincinnatiensis* (NCBI txid1123491), *Vibrio metschnikovii* (*NCBI* txid28172), *Vibrio diazotrophicus* (NCBI txid913829), *Vibrio cidicii* (*NCBI* txid1763883), *Vibrio xiamenensis* (*NCBI* txid861298), *Vibrio caribbeanicus* (*NCBI* txid701175), *Vibrio nereis* (*NCBI* txid693), *Vibrio sonorensis* (*NCBI* txid1004316), *Vibrio sinaloensis* (*NCBI* txid379097).

### MshA-GST and GST purification

For MshA-GST protein expression, an overnight culture of *E. coli* BL21 (DE3) carrying the plasmid pNUT2408 was grown at 28 °C under shaking conditions in the presence of 0.5% lactose. For GST expression, the same conditions were used for *E. coli* BL21 (DE3) carrying the plasmid pNUT2561.

Cells were collected by centrifugation and washed in PBS buffer (10 mM Na_2_HPO_4_, 1.8 mM KH_2_PO_4_, 2.7 mM KCl, 140 mM NaCl, pH 7.3), resulting in 5 mL buffer per gram of cell pellet. Next, cells were centrifuged and resuspended in PBS buffer, followed by lysis *via* sonication on ice. Next, the lysate was centrifuged twice (15,000 g, 30 min) to remove cell debris and the supernatant was collected and mixed with Protino Glutathione Agarose 4B (Macherey-Nagel # 745500.10) equilibrated with PBS buffer. The mixture was incubated end-over-end for 1 h at 4 °C then packed into a column. The column was washed extensively with PBS to remove non-specifically bound proteins. Next, the protein was eluted from the column with elution buffer (50 mM Tris-base, 10 mM glutathione, pH 8.0) and stored at −80 °C until further use.

### Biofilm matrix isolation

A Petri dish filled with 10 mL of LB broth was inoculated with *V. cholerae* C6706 wild type (KDV201). After 48 h of static incubation at 25 °C, the planktonic cells were removed from the suspension by gentle pipetting. The remaining surface-attached biofilm was washed twice by the addition of PBS, agitation on a rotary shaker for 5 min with PBS, removal of PBS and non-attached cells, and the addition of fresh PBS. The presence of surface-attached biofilms was confirmed using brightfield microscopy. These biofilms were scraped off the Petri dish using a cell scraper and matrix proteins were then prepared as described earlier (25). To separate cells from biofilm matrix, the biofilm was disrupted by vortexing for 30 min in the presence of 0.5-0.75 mm glass beads (#397641000, Acros Organics) and centrifuged to remove particulates. The final supernatant containing matrix proteins was passed through a 0.22 μm filter to obtain a clear supernatant. The freshly prepared matrix protein suspension was immediately used for the pull-down assay and for mass spectrometry.

### Identification of MshA binding partners

To identify binding partners of the MshA protein, a pull-down assay was performed (73), as illustrated schematically in Fig. S5. For this assay, the suspension of *V. cholerae* biofilm matrix proteins (prepared as described above) was equally divided into four 1.5 mL Eppendorf tubes. These tubes were mixed with either purified MshA-GST (0.04 μM, for 2 replica tubes) or with purified GST (0.04 μM, for 2 replica tubes). These mixtures were incubated for 1 h with mild rotation at 4 °C to achieve thorough mixing of bait proteins with their potential prey proteins. Then, 100 μL of GST-trap agarose resin (#sta-20, Chromotek, consisting of GST nanobody/V_H_H coupled to agarose beads) was added to the mixture, to capture GST-tagged proteins and incubated for 1 h at 4 °C on a rotary shaker. To elute the proteins from the resin, 0.5% SDS in Tris buffer (pH 8.0) was added and samples were boiled at 90 °C for 5 min. Eluted proteins were stored at −80 °C until further use.

Mass spectrometry was performed to identify the eluted proteins. Beads were washed 5x with 100 mM ammoniumbicarbonate buffer to remove detergents and protease inhibitors. For each sample, on-bead tryptic digest was carried out by adding 100 μL of a buffer containing 100 mM ammoniumbicarbonate and 1 μg trypsin to the beads for 45 min. Predigested proteins were separated from the beads and digestion was continued overnight at 30 °C. After completion of the proteolysis, the peptides were purified using C18 soli phase extraction, described in detail previously (74). Liquid chromatography-mass spectrometry (LC-MS/MS) analysis of the peptide samples was carried out on a Q-Exactive Plus instrument connected to an Ultimate 3000 RSLC nano and a nanospray flex ion source (all Thermo Fischer Scientific). Peptide separation was performed on a reverse-phase high-performance liquid chromatography column (75 μm × 42 cm) packed in-house with C18 resin (2.4 μm, Dr. Maisch). The peptides were loaded onto a PepMap 100 pre-column (Thermo Fischer Scientific) and eluted by a linear acetonitrile gradient from 2–35% solvent (99.85% ACN in 0.15% formic acid) over 60 min. The flow rate was set to 300 nl min^−1^. The peptides were analyzed in positive ion mode. The spray voltage was set to 2.5 kV, and the temperature of the heated capillary was set to 300 °C. Survey full-scan MS spectra (*m*/*z* = 375–1500) were acquired in the Orbitrap with a resolution of 70,000 full width at half maximum at a theoretical *m*/*z* 200 after accumulation a maximum of 3 × 10^6^ ions in the Orbitrap. Based on the survey scan, up to 10 most intense ions were subjected to fragmentation using high collision dissociation at 27% normalized collision energy. Fragment spectra were acquired at 17,500 resolution. The ion accumulation time was set to 50 ms for both MS survey and tandem MS (MS/MS) scans. To increase the efficiency of MS/MS attempts, the charged state screening modus was enabled to exclude unassigned and singly charged ions. The dynamic exclusion duration was set to 30 s. For label-free quantification, the MS raw data was analyzed using MaxQuant version 1.6.10.43 (75), and a UniProt protein database of *V. cholerae.* Further data evaluation was performed within Perseus (76).

### Microscopy

To obtain high-resolution images at cellular resolution of flow chamber biofilms, a Nikon Ti-E inverted microscope was used together with a 100x silicon oil immersion objective with numerical aperture 1.35 (Olympus) and a Yokogawa spinning disk confocal unit. Fluorescence was excited with a 488 nm laser (for sfGFP), a 552 nm laser (for mRuby3), or a 637nm laser (for Alexa Fluor 647). The microscope and camera were controlled with the NIS Elements Advanced Research software (Nikon).

For low-resolution imaging to quantify biofilm biovolume during dispersal, a 40x air objective with numerical aperture 0.6 (Nikon) and a 488 nm laser (for sfGFP) was used on the same microscope and confocal unit as described above.

To visualize and quantify matrix proteins inside biofilms, biofilms of *V. cholerae* strains expressing 6xHis- or HA-tagged versions of the matrix proteins were grown in our standard flow chambers as described above. These tags were attached to the native chromosomal locus of the matrix protein of interest, resulting in the strains KDV2151, KDV2186 and KDV2192. During the entire growth period of the biofilm, the inflowing medium contained M9 medium supplemented with either a anti-6xHis antibody conjugated to Alexa Fluor 555 (#MA1-135-A555, Invitrogen), or a anti-HA antibody conjugated to Alexa Fluor 647 (#26183-A647, Invitrogen) to stain the matrix proteins. To visualize and quantify the VPS polysaccharide in the biofilm matrix, biofilms were grown in M9 medium containing 20 μg/mL of wheat-germ agglutinin (WGA) conjugated to FITC and 20 μg/mL Concanavalin A conjugated to FITC (Sigma-Aldrich), and 0.5 mg/mL of filter-sterilized bovine serum albumin (BSA).

To visualize MSHA-pili inside *V. cholerae* biofilms, a derivative of the WT strain C6706 was constructed that harbors a C-terminal 6xHis-tagged *mshA* at the native locus (strain KDV2445). For the detection of MSHA in live cells, biofilms were grown for 18 h in flow chambers with M9 medium, followed by the addition of 0.5 mg/mL BSA and anti-6xHis antibody conjugated to Alexa Fluor 555 (#MA1-135-A555, Invitrogen, final concentration 2 μl/mL) for 1 h, at which point the biofilms were imaged using spinning disk confocal microscopy and the 100x silicon oil immersion objective as described above. BSA was added to the media at a final concentration of 0.5 mg/mL to avoid signal from non-specific binding of antibodies.

### Image analysis

All image analysis was performed with the software tool BiofilmQ (https://drescherlab.org/data/biofilmQ/) (61) using the cube segmentation and analysis method, which enables cytometric analysis of both low-resolution and high-resolution images of microbial communities (77). For high-resolution images and single-cell segmentation, we used an edge-detection-based segmentation approach (41) followed by a BiofilmQ analysis of the single cell properties.

For quantification of biofilm biovolume changes, fluorescence images were first registered and manual cropping was applied. Cropped biofilms were segmented using cube segmentation and global biofilm parameters were calculated. Biofilm biovolumes for each biofilm at each acquired timepoint were exported for further analysis in Matlab. For quantification of extracellular biofilm matrix, fluorescence images were first registered and biofilms cropped manually. Cells were segmented using threshold-based cube segmentation. Mean intensity of immunolabeled matrix was calculated in a 3-voxel range surrounding each segmented cell. Kymographs were plotted with cubes and their respective matrix intensity shell as function of distance from biofilm surface. For quantification of MSHA pili abundance, fluorescence images where first registered and biofilms cropped manually. Cells were segmented using 3D edge-detection and 3D watershed with an optimized seed selection. Mean intensity of immunolabeled MSHA pili was calculated in a shell of 3-voxel range surrounding each segmented cell. Kymographs were plotted with cells and their respective MshA intensity shell as a function of distance from the biofilm surface. Kymographs of biofilm matrix and MSHA pili abundance of all analyzed positions of all replicates were averaged.

### Code and data availability

The biofilm image analysis software tool BiofilmQ (61) is available online (https://drescherlab.org/data/biofilmQ/).

